# Many unreported crop pests and pathogens are probably already present

**DOI:** 10.1101/519223

**Authors:** Daniel P. Bebber, Elsa Field, Gui Heng, Peter Mortimer, Timothy Holmes, Sarah J. Gurr

## Abstract

Biotic invasions threaten global biodiversity and ecosystem function. Such incursions present challenges to agriculture where invasive pest species cause significant production losses require major economic investment to control and can cause significant production losses. Pest Risk Analysis (PRA) is key to prioritizing agricultural biosecurity efforts, but is hampered by incomplete knowledge of current crop pest and pathogen distributions. Here we develop predictive models of current pest distributions and test these models using new observations at sub-national resolution. We apply generalized linear models (GLM) to estimate presence probabilities for 1739 crop pests in the CABI pest distribution database. We test model predictions for 100 unobserved pest occurrences in the People’s Republic of China (PRC), against observations of these pests abstracted from the Chinese literature. This resource has hitherto been omitted from databases on global pest distributions. Finally, we predict occurrences of all unobserved pests globally. Presence probability increases with host presence, presence in neighbouring regions, per capita GDP, and global prevalence. Presence probability decreases with mean distance from coast and known host number per pest. The models were good predictors of pest presence in Provinces of the PRC, with area under the ROC curve (AUC) values of 0.75 – 0.76. Large numbers of currently unobserved, but probably present pests (defined here as unreported pests with a predicted presence probability > 0.75), are predicted in China, India, southern Brazil and some countries of the former USSR. Our results shows that GLMs can predict presences of pseudo-absent pests at sub-national resolution. The Chinese scientific literature has been largely inaccessible to Western academia but contains important information that can support PRA. Prior studies have often assumed that unreported pests in a global distribution database represents a true absence. Our analysis provides a method for quantifying pseudo-absences to enable improved PRA and species distribution modelling.

## Introduction

The spread of invasive species is homogenizing the biosphere, with wide-ranging implications for natural ecosystems (Baiser *et al.*, 2012; Santini *et al.*, 2013) and agriculture (Fisher *et al.*, 2012; Bebber *et al.*, 2014a; Bebber, 2015). The number of first observations of crop pests and pathogens has accelerated in recent years, driven primarily by global trade (Ding *et al.*, 2008; Bacon *et al.*, 2014), but also potentially by climate change and our improving ability to monitor and identify threats (Bebber *et al.*, 2014a; Bebber, 2015). Here, we use the term ‘pest’ to describe any herbivorous arthropod, pathogenic microbe or virus known to attack agricultural crops. Emerging pests can be extremely damaging to agricultural production and the economy, causing both pre-harvest and post-harvest losses (Bebber & Gurr, 2015; Paini *et al.*, 2016; Savary *et al.*, 2017). Recently, for example, sub-Saharan Africa has suffered from the virulent Ug99 strain of the wheat stem rust fungus (*Puccinia graminis tritici*) (Patpour *et al.*, 2015), the newly-evolved Maize Lethal Necrosis viral syndrome (Wangai *et al.*, 2012), arrival of the Asian citrus psyllid (*Diaphorina citri*) which vectors citrus greening disease (Shimwela *et al.*, 2016), and the appearance of Tropical Race 4 of *Fusarium oxysporum f. sp. cubense* attacking Cavendish bananas (Ordonez *et al.*, 2015). Central America, Europe, East Africa and Australia have been identified as hotspots of new pest invasions, with maize, bananas, citrus and potato as the crops most likely to be affected (Bebber, 2015). Outbreaks of resident pests due to favourable weather conditions, virulence evolution, or crop management factors, add to the burden on farmers. For example, a major outbreak of coffee leaf rust (*Hemileia vastatrix*) in Latin America, likely to have been triggered by a failure in disease management, is reported to have caused large-scale unemployment and social upheaval in recent years (Avelino *et al.*, 2015).

Despite the expanding ranges of many pests, complete occupation of their potential ranges has not yet occurred (Bebber *et al.*, 2014a) and so there remains a strong impetus for implementation of biosecurity measures at international borders (Fears *et al.*, 2014; Flood & Day, 2016; MacLeod *et al.*, 2016). Control of spread within countries is extremely difficult because of unhindered transport of plants and soils (Ward, 2016), and biosecurity measures focus on quarantine and inspections at borders (MacLeod *et al.*, 2016). A key component of international phytosanitary action is Pest Risk Analysis (PRA), a suite of methods that allow countries to prioritize protective measures against those pests most likely to arrive and cause serious economic damage (Robinet *et al.*, 2012; Baker *et al.*, 2014). PRA involves assessment of the likelihood of pest arrival, the likelihood of establishment, the potential economic impact if uncontrolled, and the prospect of successful control or eradication (Baker *et al.*, 2014). To date, PRA has primarily been based upon expert opinion regarding the likelihood of arrival and potential impact of individual pests. For example, the UK’s recently-established Plant Health Risk Register (PHRR) (Baker *et al.*, 2014) employs simple climate-matching (based on known pest distributions) and host availability to assign qualitative risks of invasion and impact, but not quantitative predictive models. Examples of registered pests include the Oleander aphid *Aphis nerii* which has been assigned very low likelihoods of arrival and establishment, and would cause negligible damage if it did, whereas the Zebra chip phytoplasma *Candidatus liberibacter solanacearum* is thought moderately likely to arrive and would have a very serious impact if it did (DEFRA, 2018).

While quantitative PRA protocols have been recently developed recently by the European Food Safety Authority (Jeger *et al.*, 2018), examples of quantitative PRA are rare in international phytosanitary legislation and practice. This contrasts with the long and vibrant history of research in predictive species distribution modelling (SDM) for pests (Elith & Leathwick, 2009; Sutherst, 2014). The geographic distributions of species are non-random, determined by their biotic environment (e.g. hosts or prey), the abiotic environment (e.g. climate, edaphic factors), and migration (dispersal to suitable habitat) (Soberón & Peterson, 2005; Soberón, 2007; Soberón & Nakamura, 2009). Thus, pest invasion risk is, in theory, quantifiable. Numerous modelling approaches are now available to predict the likely distributions and impacts of pests (Elith & Leathwick, 2009; Venette *et al.*, 2010; Robinet *et al.*, 2012), ranging from process-based, or mechanistic models, to statistical, or correlative approaches (Dormann *et al.*, 2012). Regional and global databases on known pest distributions are commonly used to parameterize these models, either providing direct estimates of pests’ ecological niches (Venette *et al.*, 2010; Kriticos, 2012), or indirectly via shared geographic ranges (Paini *et al.*, 2010, 2016; Eschen *et al.*, 2014).

One seldom-acknowledged issue with pest distribution data in global databases is geographic bias in the likelihood that a pest will be detected, correctly identified, reported and recorded (Pyšek *et al.*, 2008). Analysis of one of the most widely studied global pest distribution databases suggests that hundreds of pests already present in many developing countries have not been reported (Bebber *et al.*, 2014b). The total number of observed pests in an administrative area (country, or administrative division for larger countries) can be largely explained by scientific capacity and agricultural production. Under a scenario of globally high scientific and technical capacity (i.e. where all countries have US-level per capita GDP and research expenditure), analysis predicts that many countries across the developing world would report hundreds more pests. This suggests that a large fraction of the *current* agricultural pest burden is unreported and unknown, and that even the best global databases suffer from severe observational bias. This has potentially serious consequences for both plant biosecurity activities and for research based upon these databases. Such observational bias may have implications for SDM methods that infer environmental tolerances from observed distributions. Scientific capacity, economic development, and the ability to detect, identify and report pests, are strongly correlated with latitude, as is climate (Bebber *et al.*, 2014b). Under-reporting of pests at low latitudes will therefore bias estimation of climate tolerances, as occurrence is underreported in warmer regions. Reducing this observational bias by strengthening pest identification efforts in the developing world is therefore critical in improving scientific understanding of pest distributions, and in PRA.

The People’s Republic of China (henceforth referred to as China) has been predicted to harbour the largest number of pests (Bebber *et al.*, 2014b). China produces the largest quantity of crops by tonnage globally, and has the greatest diversity of production. Both factors are strong determinants of recorded pest numbers (Bebber *et al.*, 2014b). Yet, the actual recorded number of pests in China is much smaller than expected (Bebber *et al.*, 2014b). For many countries, under-reporting of agricultural pests is likely to be purely a function of the lack of institutional capacity to detect, identify, and report incidences in the scientific and ‘grey’ literature used by CABI to populate the distribution database. For China, there is potentially an interesting alternative. The Chinese literature was, until the reforms of 1978, largely inaccessible to Western academia. Even post-reform and the opening of China instigated by Deng Xiaoping, Chinese-language publications are not commonly accessed by English-speaking researchers. A scientifically-important translation of the Chinese literature is the reporting of the anti-malarial compound artemisinin (Klayman, 1985). The Chinese research literature, having developed isolated from the Western literature, therefore provides a potentially independent data source for testing models of pest distributions.

Here, we develop statistical models of global pest presence using a database of known pest occurrences and confront the predictions of individual pest or pathogen presence in China’s provinces with observations from the Chinese literature. In addition, we compare models in which pest absences are treated as true absences with models in which absences are weighted according to estimates of scientific and technical capacity of a given country to report plant health risks, to investigate the effect of observational bias and pseudoabsences in pest distribution modelling. We then apply our distribution models globally to all unreported pests in all regions, to give predicted probabilities of presence. Finally, we list those pests that are probably present, but as yet unreported, around the world.

## Materials and Methods

We obtained pest distribution data from the CABI Knowledge Bank database in January 2014 with permission (CABI, Wallingford, UK). The database comprised 91,030 records of the observed distributions of 1901 agricultural pests by administrative division of each country, e.g. US States. In total, 384 geographical units were included in the model, comprising 221 countries plus sub-national divisions for Australia (7), Brazil (28), Canada (13), China (31), India (33), and the USA (51). Geographical regions such as Bouvet Island which were smaller than a single pixel (5 arc minute resolution, or approximately 100 sq. km) of the gridded crop distribution database we employed were excluded from the analysis. Host crop spatial distributions for 175 crops at 5 arc minute resolution were obtained from the EarthStat database (http://www.earthstat.org/; Monfreda *et al.*, 2008). Known plant hosts of each pest were taken from the CABI Knowledge Bank, and the host genera matched to the genera in the list of 175 crops. Pests without known hosts in this list of 175 crops were excluded from the analysis. Pests from taxonomic groups with fewer than 50 species (e.g. Acari, Gastropoda and various other insect taxa) were also excluded from the analysis. This left a total of 1739 pests comprising 124 species, subspecies and pathotypes of Bacteria, 106 Diptera, 215 Coleoptera, 398 Fungi, 233 Hemiptera, 248 Lepidoptera, 99 Nematoda, 61 Oomycota and 209 viruses. Assigning reported presences for each pest to each geographical region gave a dataset of 667,776 presences or absences for each pest-region pair. In total, there were 81,821 presences (12.2 per cent of the total) in the final dataset.

We developed Generalized Linear Models (GLM), using the *glm* function (*MASS* package) in R v.3.4.0 with logit link for binomial data (R Development Core Team, 2017), for the presence or (pseudo-) absence of each pest in each region. Model predictors were as follows: log-transformed *per capita* GDP for the country as a whole in 2016 (World Bank data, http://data.worldbank.org/); log-transformed total number of crop host genera for the pest (CABI Knowledge Bank, obtained with permission); log-transformed area (ha) of the pest’s host crop distribution (summing planted areas of all known host crops in each geographical region); log-transformed host crop area (ha) of neighbouring (i.e. with land border) regions which have reported the pest as present (set to zero if no neighbours have reported the pest); log-transformed total fraction of regions globally that have reported the pest; and log transformed distance (km) of crop area to the coast (calculated as the distance of the centroid of the crop area distribution from the nearest coastline). Log transformations were applied to distribute the predictor variable values more evenly across the sample space. Briefly, the rationale for these predictors was that GDP is a proxy for historical trade (Pyšek *et al.*, 2010) and observational capacity (Bebber *et al.*, 2014b), host area indicates the available habitat for each pest, host number indicates the degree of biotic generalism of the pest, neighbouring-region presence indicates the potential for spread across a land border, fraction of regions reporting presence indicates global ubiquity and environmental generalism, and distance to coast indicates proximity to international shipping ports (Chapman *et al.*, 2017).

We developed two pest distribution models. The ‘unweighted’ model included geographical and biological predictors and treated all unobserved pests as absent from a region. The ‘weighted’ model treated unobserved pests as potentially pseudo-absent, using a function of the scientific and technical capacity of each country (Bebber *et al.*, 2014b). Presences were taken as being correct and unambiguous, and given a weighting of unity. Absences were weighted by the logarithm of the agricultural and biological sciences publication output of each country from 1996 – 2016 (Scimago Lab, 2017), normalized to the logarithm of the output of the USA (the world’s most scientifically productive country), such that the absence weight *w*_0_ = log(*s*)/log(*s*_USA_). Thus, pests unreported from scientifically advanced nations were assumed not to be present (or, present at undetectable population density), while pests unreported from developing nations were less informative of absence. China, with the second largest research output, had *w*_0_ = 0.93, suggesting that non-reporting of a pest should be relatively strong evidence of its physical absence. However, we hypothesized that non-reporting in the CABI databases could be due to lack of translation from the Chinese literature, therefore we set *w*_0_ to zero for China, effectively omitting these pseudo-absences from the analysis. The weighted and unweighted models were compared with a null model assuming constant presence probability using Likelihood Ratio Tests.

To validate the models we predicted the probability of presence for a random sample of 100 as-yet unobserved pests in all Chinese Provinces, but excluding Taiwan. The Chinese literature was searched for observations of these unobserved pests in China. We used the text mining methodology designed by CABI for their Plantwise Knowledge Bank. The following rules were followed to locate pest records in the Chinese literature:

- Include only papers that are primarily about distribution data, not those where distribution is mentioned, but where something else is the primary focus. If this is unclear do not process the paper.
- Mine only the primary literature (including Masters and Doctoral theses), not meta-analyses, reviews, or non-peer reviewed (“grey”) literature.
- Pest and host names must be preferred scientific names, following the CAB Thesaurus (www.cabi.org/cabthesaurus/) and the Plant List (http://www.theplantlist.org/).
- Record country and location information given in the paper, including latitude/longitude. CABI uses five levels for location, from the largest scale (i.e., provincial) to the smallest (i.e., village/town).
- Record date of observation/collection (entering each year separately) and date of publication. Can be left blank if not given, or use the date of receipt in the diagnostic laboratory as a surrogate for date of collection.
- Record pest status – present/not found. Only record absence if pest absence is specifically stated in the paper.
- Record pest status using only the status terms defined by CABI, and only if used in the paper e.g. “widespread”, “restricted” “soil only” “greenhouse only” (see CABI guidelines for complete list).
- Record if the paper was a first record of that pest or not and details of this (e.g. “first record in <country/location>”, “first record on <host species name>”)
- Only enter data where the pest/pathogen has been clearly identified, not just symptoms seen.
- Record only natural infections, not artificial inoculants.

Combinations of pests and locations were submitted to several search engines. The priority of search engines was: Baidu (www.baidu.com), China National Knowledge Infrastructure (CNKI, http://www.cnki.net), Chongqing VIP Information Company (CQVIP, http://lib.cqvip.com/), and Wangfang Data (http://www.wanfangdata.com.cn). Baidu is the most popular Chinese internet search engine. CNKI is led by Tsinghua University, and supported by ministries of the Chinese Government. CQVIP, formerly known as Database Research Center under the Chongqing Branch of the Institute of Scientific & Technical Information of China (CB-ISTIC), was China’s first Chinese journal database research institution. Wanfang Data is an affiliate of the Chinese Ministry of Science & Technology, and provides access to a wide range of database resources.

Publication titles were searched first, followed by full text interrogation. The first 50 search results were scanned before dismissing a search term. The first search term combination was pest name and location (Province). If this yielded no result, then pest name and various distribution terms were tried. These distribution terms were: “catalogues” OR “checklists” OR “distribution” OR “inventories” OR “new records” OR “surveys” OR “geographical distribution” OR “new geographic records” OR “new host records”. Searches included local names in Chinese where these were known or could be identified from the literature, preferred scientific names, and non-preferred scientific names from CAB Thesaurus (https://www.cabi.org/cabthesaurus/). Searches continued until one piece of literature was found for that pest in that region, that fitted all of the requirements for CABI text mining.

If a pest was not found in any of these searches, it was assumed to be absent from the literature and thus effectively absent from the region. We cannot prove, however, that a pest is present at very low population density and has not yet been detected (Crooks, 2005).

Modelled probabilities of reported pest presence in the global dataset, *P*_*G*_, were obtained from the predictor variables for each pest-region combination, for each GLM (*predict* function in R). We then compared *P*_*G*_ with the observed presence-absence data for our Chinese sample data using logistic regressions (*glm* function in R) and Receiver-Operator Characteristic (ROC) curves (*pROC* library for R). The logistic regression coefficients *c* and *m* determine the probability of pest presence in the Chinese sample as *P*_*C*_ = 1/(1 + exp(-(*c* + *mP*_*G*_))). ROC curves describe the relationship between the true positive rate (sensitivity, the fraction of presences correctly identified as presences) and false positive rate (1 – specificity, where specificity is the fraction of absences correctly classified as absences) as the threshold for a binary classifier is decreased from one (classifying any presence probability less than one as absent) to zero (classifying any positive probability as present). A good predictor will have a high true positive rate and low false positive rate for any classification threshold, whereas a poor predictor will have roughly equal true and false positive rates (i.e., be uninformative). The Area Under Curve (AUC) for the ROC curves gives the probability that, for a random pair of presence and absence observations, the presence probability will be greater for the presence than the absence (Jiménez-Valverde, 2012). Models with good discrimination ability should have AUC significantly greater than half.

For illustration, we identified probably present pests (PPP) as those for which are currently unreported from a particular region, but for which *P*_*G*_ > 0.75 in our weighted model. This threshold was chosen based on the Kent scale which suggests a probability of 0.75 as an event that would generally be described as ‘probable’ (Kent, 1994). This is an arbitrary definition but allows us to suggest some of the pests that PRA and phytosanitary activities may want to focus on.

## Results

Globally, *P*_*G*_ increased significantly with presence in neighbouring regions, the area of host crops, the global prevalence of the pest and per capita GDP in both models (Table 1). *P*_*G*_ declined with mean distance from the coast and known host crop genera per pest. The models explained similar fractions of the deviance, and had very similar ROC curves with AUC around 88 per cent (Table 1). *P*_*G*_ was always higher for the weighted model, because absences were down-weighted (i.e. fewer true zeros), but predictions for the two models were very highly correlated (r = 0.98). The models found the highest *P*_*G*_ for Hemiptera and Lepidoptera, and lowest for Nematoda, Bacteria and Acari, compared with other taxonomic groupings.

**Table 1.** GLMs for global pest presence. The unweighted model treated unobserved pests as true absences. The weighted model weighted pseudo-absences as a function of country scientific capacity. The unweighted model had AIC = 339872, AUC = 0.88, Nagelkerke R^2^ = 0.40, McFadden R^2^ = 0.32. The weighted model had AIC = 308171, AUC = 0.88, Nagelkerke R^2^ = 0.37, McFadden R^2^ = 0.31. CoastDist is distance of crop centroid from the coast (km), GDP is per capita GDP (US$), Hosts is reported number of host crop genera, HostArea is harvested area of known host crops, NeigArea is harvested area of host crops in neighbouring regions that have reported the pest, and Prevalence is the fraction of all regions that have reported the pest.

|  | Unweighted model |  |  |  | Weighted model |  |  |  |
| --- | --- | --- | --- | --- | --- | --- | --- | --- |
|  | Mean | SE | Z | Pr(>\|Z\|) | Mean | SE | Z | Pr(>\|Z\|) |
| Acari (Intercept) | -3.67 | 0.051 | -72.3 | 0.000 | -0.897 | 0.055 | -16.3 | 0.000 |
| + Bacteria | -0.091 | 0.032 | -2.9 | 0.004 | -0.073 | 0.034 | -2.2 | 0.014 |
| + Coleoptera | 0.036 | 0.030 | 1.2 | 0.240 | 0.039 | 0.032 | 1.2 | 0.180 |
| + Diptera | 0.092 | 0.034 | 2.7 | 0.006 | 0.104 | 0.036 | 2.9 | 0.026 |
| + Fungi | 0.027 | 0.028 | 1.0 | 0.337 | 0.033 | 0.030 | 1.1 | 0.380 |
| + Hemiptera | 0.167 | 0.029 | 5.7 | 0.000 | 0.150 | 0.031 | 4.8 | 0.000 |
| + Lepidoptera | 0.145 | 0.029 | 4.9 | 0.000 | 0.134 | 0.032 | 4.2 | 0.000 |
| + Nematoda | -0.150 | 0.033 | -4.5 | 0.000 | -0.143 | 0.035 | -4.1 | 0.000 |
| + Oomycota | 0.046 | 0.034 | 1.4 | 0.176 | 0.061 | 0.037 | 1.7 | 0.151 |
| + Virus | 0.047 | 0.030 | 1.6 | 0.120 | 0.067 | 0.033 | 2.1 | 0.137 |
| log(CoastDist + 1) | -0.176 | 0.004 | -49.0 | 0.000 | -0.222 | 0.004 | -57.7 | 0.000 |
| log(GDP + 1) | 0.295 | 0.004 | 81.5 | 0.000 | 0.086 | 0.004 | 22.3 | 0.000 |
| log(Hosts + 1) | -0.300 | 0.004 | -71.4 | 0.000 | -0.297 | 0.005 | -65.6 | 0.000 |
| log(HostArea + 1) | 0.171 | 0.001 | 123.1 | 0.000 | 0.159 | 0.001 | 108.1 | 0.000 |
| log(NeigArea + 1) | 0.140 | 0.001 | 181.2 | 0.000 | 0.142 | 0.001 | 173.5 | 0.000 |
| log(Prevalence) | 0.842 | 0.007 | 124.3 | 0.000 | 0.867 | 0.007 | 121.3 | 0.000 |

We validated the models with reports of pests abstracted from the Chinese literature.

For illustration, we defined a ‘probably present pest’ (PPP) as one unreported from a region, but with *P*_*G*_ > 0.75 (using the weighted model). Overall, only 4702 of 585955 (0.8 per cent) of all unreported pest-region combinations fell into this class (Supplementary Table S1). The number of PPPs per pest category was greatest for Fungi (2052) and Hemiptera (859). Overall, 86 per cent of unreported pest-region combinations were predicted to be unlikely (*P*_*G*_ < 0.25). China, India, the USA and Eastern Europe had the largest numbers of predicted PPPs, along with other parts of East Asia and Southern Brazil (Figure 1). The top ten PPPs by number of global regions were *Cochliobolus heterostrophus* (Ascomycota: Pleosporales, a pathogen of maize), *Rhopalosiphum padi* (Arthropoda: Hemiptera, cereal pest), *Gibberella fujikuroi* (Ascomycota: Hypocreales, rice pathogen), *Sitophilus zeamais* (Arthropoda: Coleoptera), maize and rice pest), *Schizaphis graminum* (Arthropoda: Hemiptera, pest of Poaceae cereals), *Setosphaeria turcica* (Ascomycota: Pleosporales, maize pathogen), *Aphis spiraecola* (Arthropoda: Hemiptera, wide host range), *Nezara viridula* (Arthropoda: Hemiptera, legume pest), *Acyrthosiphon pisum* (Arthropoda: Hemiptera, legume pest) and *Rhopalosiphum maidis* (Arthropoda: Hemiptera, pest of maize and other crops).

**Figure 1.**
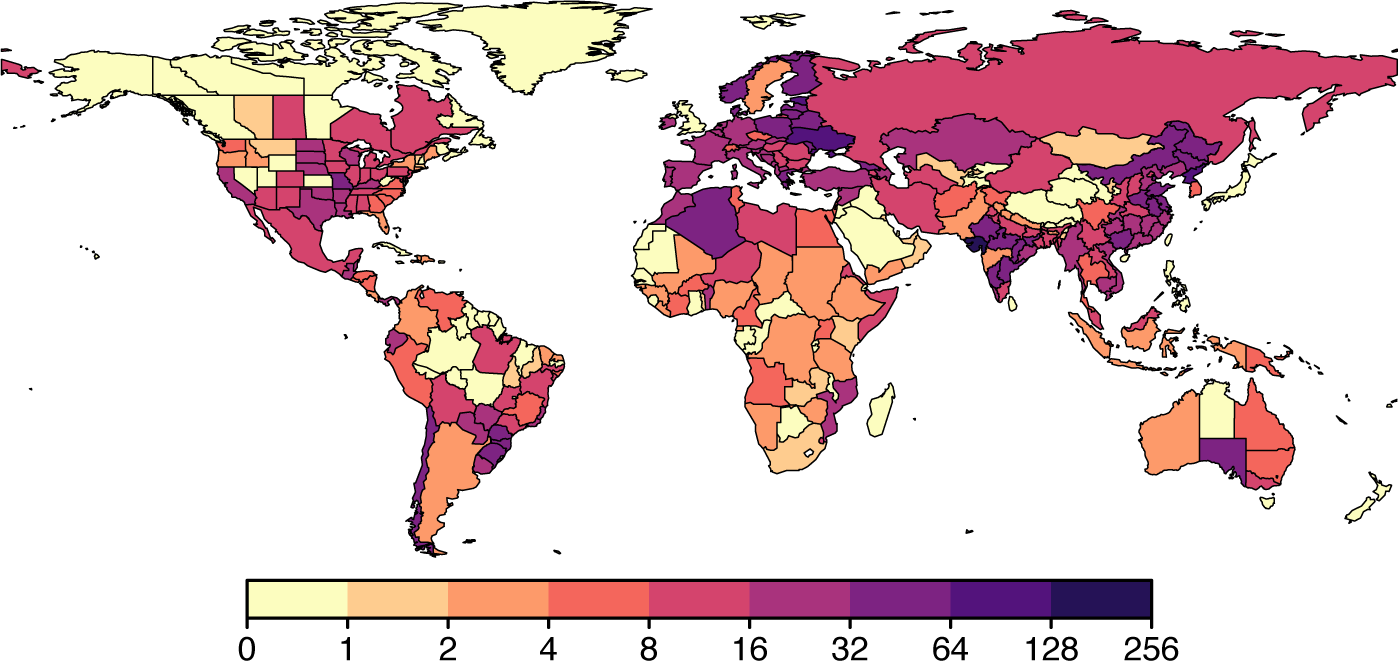
Total number of probably present pests (PPP) in all countries and sub-national regions.

Total numbers of recorded pests in China’s Provinces and municipalities increased from northern and central regions to southern and coast regions (Figure 2a), except for the central province of Gansu which had 826 reported pests. There is no obvious reason why numbers would be so large in Gansu. Here, agricultural production is moderate, and there are no particular academic centres which could account for observational bias. Hence, the Gansu values appear to be an artefact of the CABI database. The smallest numbers of recorded pests were in the mountainous provinces of Qinghai (0) and Xizang Zizhiqu (Tibet) (73), the central provinces of Ningxia (48), and the municipalities of Chongqing (24), Tianjin (3), Beijing (50) and Shanghai (55). Total numbers were largest in the coastal provinces of Guangdong (301), Zeijiang (294), Jiangsu (293), Fujian (263), and also in the southern provinces of Yunnan (291) and Sichuan (259).

**Figure 2.**
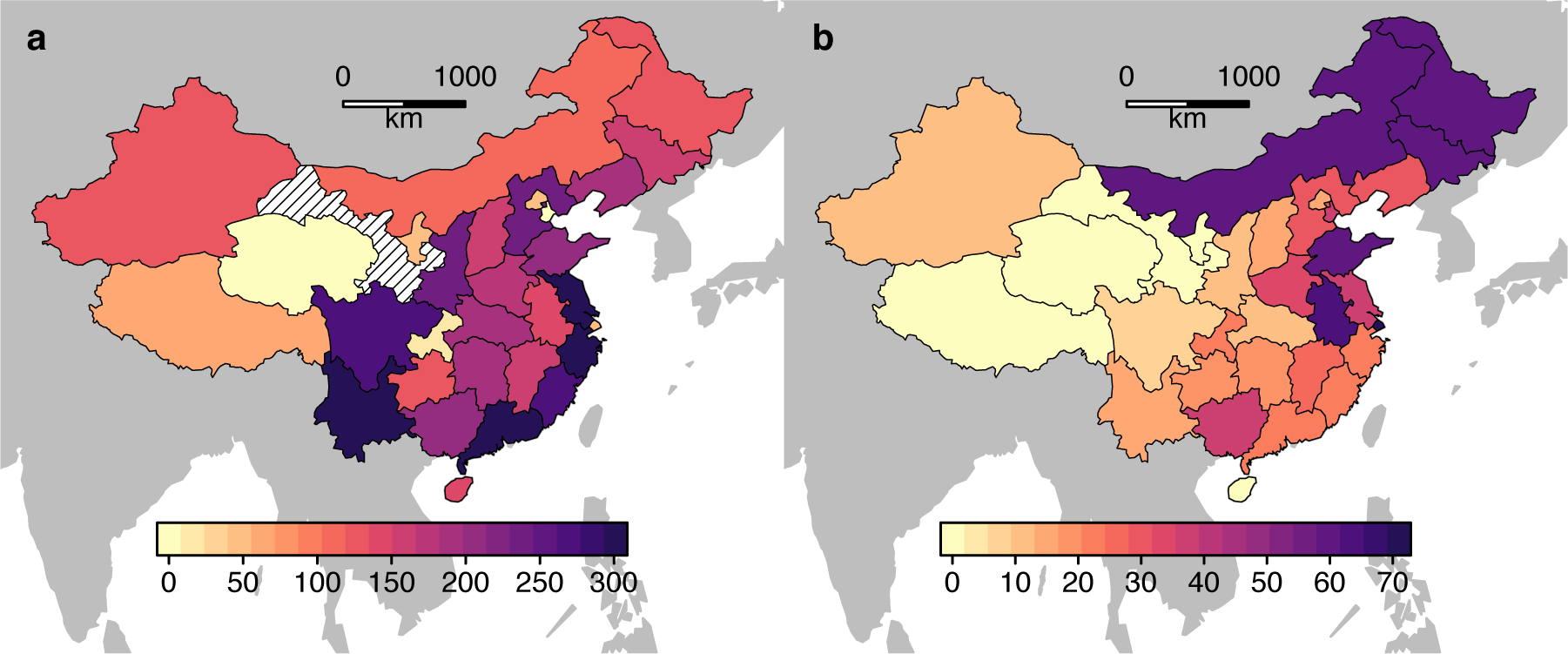
a) Total number of pests recorded in the CABI pest distribution database by China Province (excluding Taiwan). Hatched region is Gansu, see text for details. b) Total number of probably present pests (PPP) in China Provinces.

We validated our models using published pest observations from the Chinese literature. Both models were significant predictors of pest presence/absence for 100 randomly-sampled pest-Province combinations, of which 27 were found to be present (Figure 3, Supplementary Table S2). For the unweighted model, the coefficients of the logistic function were *c* = −1.73 ± 0.34 and *m* = 3.52 ± 1.25 (likelihood ratio test *vs* null model, p = 0.0043). For the weighted model, the coefficients were −1.90 ± 0.38 and 3.19 ± 0.96 (likelihood ratio test, p = 0.0006). The predictive power of the models was also tested using ROC curves, demonstrating significant discriminant ability with AUC of 0.76 (95 per cent Confidence Interval 0.66 – 0.86) for the unweighted model, and AUC 0.75 (0.64 – 0.86) for the weighted model (Figure 3). Our analysis revealed gaps in the CABI database, which is commonly used for analyses of global pest distributions. Taking one important potato pest, late blight *Phytophthora infestans* (Oomycota), as an example, high presence probabilities (> 0.75) were predicted for ten provinces listed as not reporting this pest in the CABI database. However, this pathogen has been reported present throughout the potato-growing regions of China, including Guangdong (Guo *et al.*, 2010).

**Figure 3.**
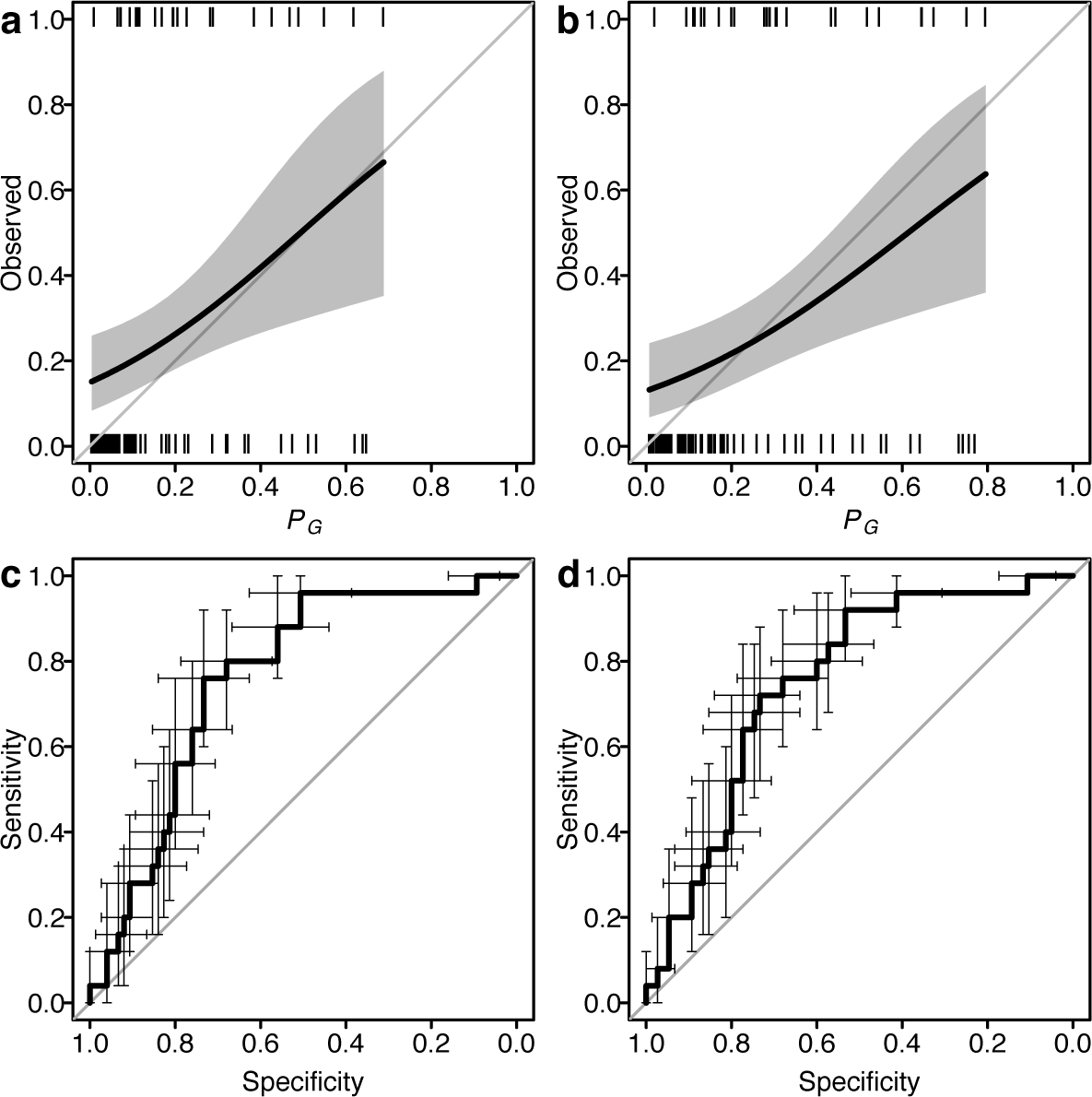
Model prediction tests. Observed presence/absence of 100 pest-province combinations vs. *P*_*G*_ from a) unweighted model and b) weighted model. Curves and shaded areas show mean and 95% CI for logistic regression fits. Tick marks show observed data. Grey diagonals show identity relationship. ROC curves for c) unweighted and d) weighted models. Error bars show 95% CI for specificity and sensitivity derived from 2000 bootstrap replications.

For China, the total number of PPPs increased from west to east (Figure 2b), and was greatest in the north eastern provinces of Jilin (59), Heilongjiang (58), and Inner Mongolia (58), the eastern provinces of Shandong (60) and Anhui (61), well as the ports of Shanghai (71) and Tianjin (51). The eastern provinces of Xizang Zizhiqu (Tibet) (1), Qinhai (1), Gansu (0) and Ningxia (2) had the lowest numbers, along with the island of Hainan (0) (Figure 3). The total number of PPPs in China was 827, the majority being Fungi (332) and Hemiptera (175). The top ten most-common PPPs in China were (in decreasing order) *Gibberella fujikuroi* (Ascomycota: Hypocreales, rice pathogen), *Aphis spiraecola* (Arthropoda: Hemiptera, generalist), *Delia platura* (Arthropoda: Diptera, pest of legumes), *Acyrthosiphon pisum* (Arthropoda: Hemiptera, legume pest), *Rhopalosiphum padi* (Arthropoda: Hemiptera, cereal pest), *Schizaphis graminum* (Arthropoda: Hemiptera, pest of Poaceae), *Curvularia* sp. (Fungi: Ascomycota, generalist pathogen), *Rhopalosiphum maidis* (Arthropoda: Hemiptera, pest of maize and other crops), *Agrotis ipsilon* (Arthropoda: Lepidoptera, generalist pest), *Lasiodiplodia theobromae* (Ascomycota: Botryosphaeriales, generalist pathogen). Thus, many of the most common PPPs in China were also common globally.

## Discussion

The Chinese literature provided strong and significant support for the predictions of pest distribution models based upon host distribution, pest prevalence, and other socioeconomic factors. China’s growing economy is expected to lead to large influxes of invasive species, including pests, in coming years (Ding *et al.*, 2008). Analysis of temporal trends in CABI pest observations show a relatively smooth increase in pests from 1950-2000, but the pattern for China is more complex, with a slow increase from 1950 until the late 1970s, a step increase, and then a more rapid growth in pest numbers from 1980 onwards (Bebber *et al.*, 2014a). One potential determinant of this sudden acceleration is the strong support for science and technology given by Deng Xiaoping in 1978, which lead to an increase in funding and academic freedom following the anti-intellectualism of the Cultural Revolution. China now ranks second only to the USA in annual R&D expenditure (IMF, 2013) and scientific output (Scimago Lab, 2017).

We identified a number of pests that were very likely to be present, and the majority of these PPPs were globally distributed and had wide host ranges. Their distributions commonly spanned wide latitudinal ranges, indicating broad climatic tolerances. *C. heterostrophus*, or Southern Leaf Spot, is primarily known as a pathogen of maize but has a wide host range. It has a wide geographic distribution both latitudinally and across continents, resulting in a high likelihood of occurrence in other regions where hosts are present. For example, *C. heterostrophus* is currently recorded only in eastern regions of North America, where most maize is grown. The lack of reported observations in the western regions of North America may be due to the fact that maize, the major host, is uncommon, and hence the disease currently has little impact. *C. sativus*, causing root and foot rot, also has a very wide geographic distribution, but an even wider host range. It is reported from Texas, Oklahoma, Mississippi, Illinois and Tennessee, but not from neighbouring Arkansas or Missouri. Hence, the high presence probability in these States. A similar pattern is seen for the maize pathogen *S. turcica.* Another global species, *R. maidis*, the green corn aphid, is reported across Europe and in Russia, but, like many other pests, not from the former Soviet states of Ukraine, Belarus, Lithuania, Latvia and Estonia. It is plausible that reporting from these nations was less likely when they were part of the USSR.

Predictors like host availability, presence in neighbouring territories and global prevalence were expected to have positive relations with presence probability. The negative relation with distance from coast is likely to be related to import via shipping ports (Huang *et al.*, 2012; Liebhold *et al.*, 2013), and supports the observation that islands report more pests than countries with land borders (Bebber *et al.*, 2014b). Detailed modelling of individual pest climate responses (Bregaglio *et al.*, 2012; Kriticos *et al.*, 2013) for such a large number of pests was beyond the scope of this study. Implicitly, we can assume that the presence of the host crop indicates that the climate is suitable for the pest (Paini *et al.*, 2016), though we acknowledge that this is not necessarily the case (Berzitis *et al.*, 2014). The negative relationship with number of host genera per pest might suggest that host specialists are more likely to invade and establish than host generalists, once host availability has been taken into account. For the practical purposes of PRA, our models provide reliable probability estimates for the presence of unreported pests at subnational resolution, and we have provided a global list of the unreported pests whose presence is most likely (Table S2).

We addressed the issue of pseudo-absences in the CABI data by statistically weighting missing pest observations in proportion to the scientific output of the reporting nation, since scientific output had been confirmed as a strong determinant of total reported pest numbers (Bebber *et al.*, 2014b). Often, unreported pests are treated as true absences in pest risk analyses (Paini *et al.*, 2016). The positive relation of GDP with presence probability supports our hypothesis that wealthy countries are more likely to detect and report pests (Bebber *et al.*, 2014b). Once observational bias is controlled for using scientific capacity-based weighting, per capita GDP becomes a weaker determinant. Our weighted model has similar overall explanatory power to our unweighted model. Nevertheless, the issue of observational biases related to country-level socioeconomic variation has been raised several times for various classes of organism (Jones *et al.*, 2008; Pyšek *et al.*, 2008; Westphal *et al.*, 2008; Boakes *et al.*, 2010; Bebber *et al.*, 2013, 2014b), and we therefore recommend the application of appropriate statistical controls when analysing datasets produced from reports of species presences (as opposed to distributional datasets derived from rigorous sampling protocols).

Our SDM was statistical, fitting response functions for various predictors to the probability of pest presence. Many SDM approaches exist, from highly mechanistic models based on pest biology and ecology (Bregaglio *et al.*, 2012; Skelsey *et al.*, 2016) to purely statistical models that utilize only patterns in known distributions (Paini *et al.*, 2010). The rarity of quantitative model input into PRAs is partly due to the scarcity of empirical data available on pest biology and epidemiology required to parameterize mechanistic models, and so key biological parameters are often inferred from known distributions (Robinet *et al.*, 2012). This is particularly the case for newly emergent pathogens for which experimental investigations have not yet been conducted. The European Food Safety Authority (EFSA) has developed quantitative PRA guidelines that recommend modelling approaches and data sources for assessing invasion and establishment risk (Jeger *et al.*, 2018), and application of these methods was attempted for *Diaporthe vaccinii*, a pest of blueberries (Jeger *et al.*, 2017). However, most of the epidemiological data required for this pest was unavailable, and the risk assessment was thus based on expert opinion or data from related pests (Jeger *et al.*, 2017). Epidemiological parameters can be poorly constrained even for long-established pests. For example, coffee leaf rust fungus (*Hemileia vastatrix*) has affected coffee production for more than a century, but a recent infection model relied upon temperature response functions derived from the single available study published three decades previously (Bebber *et al.*, 2016). Initiatives such as the EU-funded PRATIQUE project (2008-11) have attempted to fill this knowledge gap and enable modelling by collating available ecophysiological data for insect pests (Baker, 2012). While the advantages and disadvantages of the many different pest distribution and impact models continue to be researched and debated (Venette *et al.*, 2010; Dormann *et al.*, 2012; Robinet *et al.*, 2012; Sutherst, 2014), it is clear that practical application of these methods in PRA remains limited.

SDM for pests has direct policy implications for PRA and plant biosecurity. PRA is guided by International Standards for Phytosanitary Measures (ISPM), which are part of the International Plant Protection Convention (IPPC) (MacLeod *et al.*, 2010). ISPMs tend to rely on expert judgement for PRA, rather than quantitative modelling to support decision making. ISPM No. 21 “Pest Risk Analysis for Regulated Non-Quarantine Pests”, endorsed in 2004, mentions use of pest and host life-cycle and epidemiological information, but not quantitative modelling (FAO, 2004). Individual PRAs similarly employ a qualitative approach. For example, the Australian Government’s PRA for *Drosophila suzukii* references only a single unpublished report on SDM for this species, conducted for North America. Probabilities of *D. suzukii* spread within Australia are qualitatively assessed by comparison with observations in other countries (Department of Agriculture, Fisheries and Forestry, 2013). The European and Mediterranean Plant Protection Organization (EPPO) PRAs occasionally include model results. For example, a climate matching for the bacterium *Xanthomonas axonopodis* pv. *allii* was undertaken using the CLIMEX model, to identify areas at risk within the EPPO region (EPPO, 2008). However, as discussed previously, appropriate empirical studies are rare (Jeger *et al.*, 2017). Our results contribute to the quantification of risk within PRA by providing probabilistic estimates for the presence of hundreds of unreported pests around the world, thereby improving understanding of the threats to global agriculture. With growing evidence that pest ranges are shifting poleward in response to global climate change (Bebber *et al.*, 2013), our poor knowledge of pest distributions, particularly in the developing world, is troubling, both because of the burden these organisms place on farmers who have little access to detection and control technologies, and because invasions of temperate regions are likely to occur from warmer regions. Improved targeting of phytosanitary measures through quantitative PRA is therefore vital to crop protection.

## Acknowledgements

We thank the British Society for Plant Pathology for an Undergraduate Vacation Bursary for Elsa Field. We thank Le Yei and Li Huili (Kunming Institute of Botany) for assistance in text mining the Chinese literature and Ho Wai Yim for translations. CABI is an international intergovernmental organisation and we gratefully acknowledge the core financial support from our member countries (and lead agencies) including the United Kingdom (Department for International Development), China (Chinese Ministry of Agriculture), Australia (Australian Centre for International Agricultural Research), Canada (Agriculture and Agri-Food Canada), Netherlands (Directorate-General for International Cooperation), and Switzerland (Swiss Agency for Development and Cooperation). See https://www.cabi.org/about-cabi/who-we-work-with/key-donors/ for full details.

## Author contribution

DB conducted the analyses and wrote the manuscript. EF and GH searched the Chinese literature. TH assisted with CABI data acquisition. All authors contributed ideas and edited the manuscript.

## Data accessibility

Pest distribution data are available with permission from CABI, Nosworthy Way, Wallingford, OX10 8DE, UK

Sources for other datasets used in the analysis are given in the text.

